# How far are we in the rapid prediction of drug resistance caused by kinase mutations?

**DOI:** 10.1101/2020.07.02.184556

**Authors:** Mehmet Erguven, Tülay Karakulak, M. Kasim Diril, Ezgi Karaca

## Abstract

Protein kinases regulate various cell signaling events in a diverse range of species through phosphorylation. The phosphorylation occurs upon transferring the terminal phosphate of an ATP molecule to a designated target residue. Due to the central role of protein kinases in proliferative pathways, point mutations occurring within or in the vicinity of ATP binding pocket can render the enzyme overactive, leading to cancer. Combatting such mutation-induced effects with the available drugs has been a challenge, since these mutations usually happen to be drug resistant. Therefore, the functional study of naturally and/or artificially occurring kinase mutations have been at the center of attention in diverse biology-related disciplines. Unfortunately, rapid experimental exploration of the impact of such mutations remains to be a challenge due to technical and economical limitations. Therefore, the availability of kinase-ligand binding affinity prediction tools is of great importance. Within this context, we have tested six state-of-the-art web-based affinity predictors (DSX-ONLINE, KDEEP, HADDOCK2.2, PDBePISA, Pose&Rank, and PRODIGY-LIG) in assessing the impact of kinase mutations with their ligand interactions. This assessment is performed on our structure-based protein kinase mutation benchmark, BINDKIN. BINDKIN contains 23 wild type-mutant pairs of kinase-small molecule complexes, together with their corresponding binding affinity data (in the form of IC_50_, K_d_, and K_i_). The web-server performances over BINDKIN show that the raw server predictions fail to produce good correlations with the experimental data. However, when we start looking in to the direction of change (whether a mutation improves/worsens the binding), we observe that over K_i_ data, DSX-ONLINE achieves a Pearson’s R correlation coefficient of 0.97. When we used homology models instead of crystal structures, this correlation drops to 0.45. These results highlight that there is still room to improve the available web-based predictors to estimate the impact of protein kinase point mutations. We present our BINDKIN benchmark and all the related results online for the sake of aiding such improvement efforts. Our files can be reached at https://github.com/CSB-KaracaLab/BINDKIN

## 1. INTRODUCTION

Protein kinases regulate various cell signaling events in a diverse range of species through phosphorylation.^1^ The phosphorylation occurs upon transferring the terminal phosphate of an ATP molecule to a designated serine/threonine/tyrosine/histidine target residue. This transfer leads to an addition of three negative charges per phosphosite, thus, to an alteration of the physicochemical properties of target protein. The phosphorylation induces critical cellular events, such as apoptosis, and transcriptional regulation. Due to their essential role in the cellular homeostasis, a deregulation in the kinase activity often results in malignancies. Such deregulations stem from expression level changes or point mutations, which are occurring at around the kinase’s ATP binding pocket.

### 1.1. A brief introduction to the protein kinase catalysis mechanism

The protein kinases mediate the catalysis through its inter-domain interactions (between the globular N-terminal and C-terminal domains) (Figure 1). This architecture grants modularity to the enzyme to leverage binding of ATP, cofactors, and other proteins during enzyme’s catalytic cycle^2^. The smaller N-terminal lobe is constituted of antiparallel β-sheets, where the bigger C-terminal lobe is enriched in α-helices.^3–5^ The protein kinases confer four catalytically important regions, i.e., the gatekeeper residue, the activation loop, the DFG motif, and the glycine-rich loop. The kinase gatekeeper residue resides within the ATP binding pocket (Figure 2). In the inactive state of the enzyme, the 20-30 residues long activation loop is found in the DFG-out conformation. In this state, the catalytic aspartate (D), responsible for transfer of the γ-phosphate from ATP to the substrate, blocks the catalytic cleft for substrate entry (Figure 1). In the active DFG-in conformation, the side chain of the catalytic aspartate faces the ATP binding pocket, making the pocket accessible to substrate binding^6,7^(Figure 1). The conformational transition from DFG-out to DFG-in states is induced upon phosphorylation of the activation loop or by binding of ATP-competitive kinase inhibitors.^8–10^ In its active form, the catalytic aspartate chelates the essential cofactor divalent cations, either manganese or magnesium^11^ (Figure 2). Together with these cations, ATP molecule is coordinated by the glycine rich loop, and a conserved lysine of the ATP binding pocket. In this coordination, the N1 and N6 nitrogen atoms of the adenine ring form specific hydrogen bonds with the backbone of the inter-domain hinge region (Figure 2).^12^ The partially conserved nonpolar aliphatic residues (leucine, valine, phenylalanine, alanine, and methionine) present within the ATP binding pocket provide van der Waals contacts with ATP’s purine moiety.^13^ The details on the reaction mechanism are provided in the Supplementary Information.^4,5,14–18^

**Figure 1.**
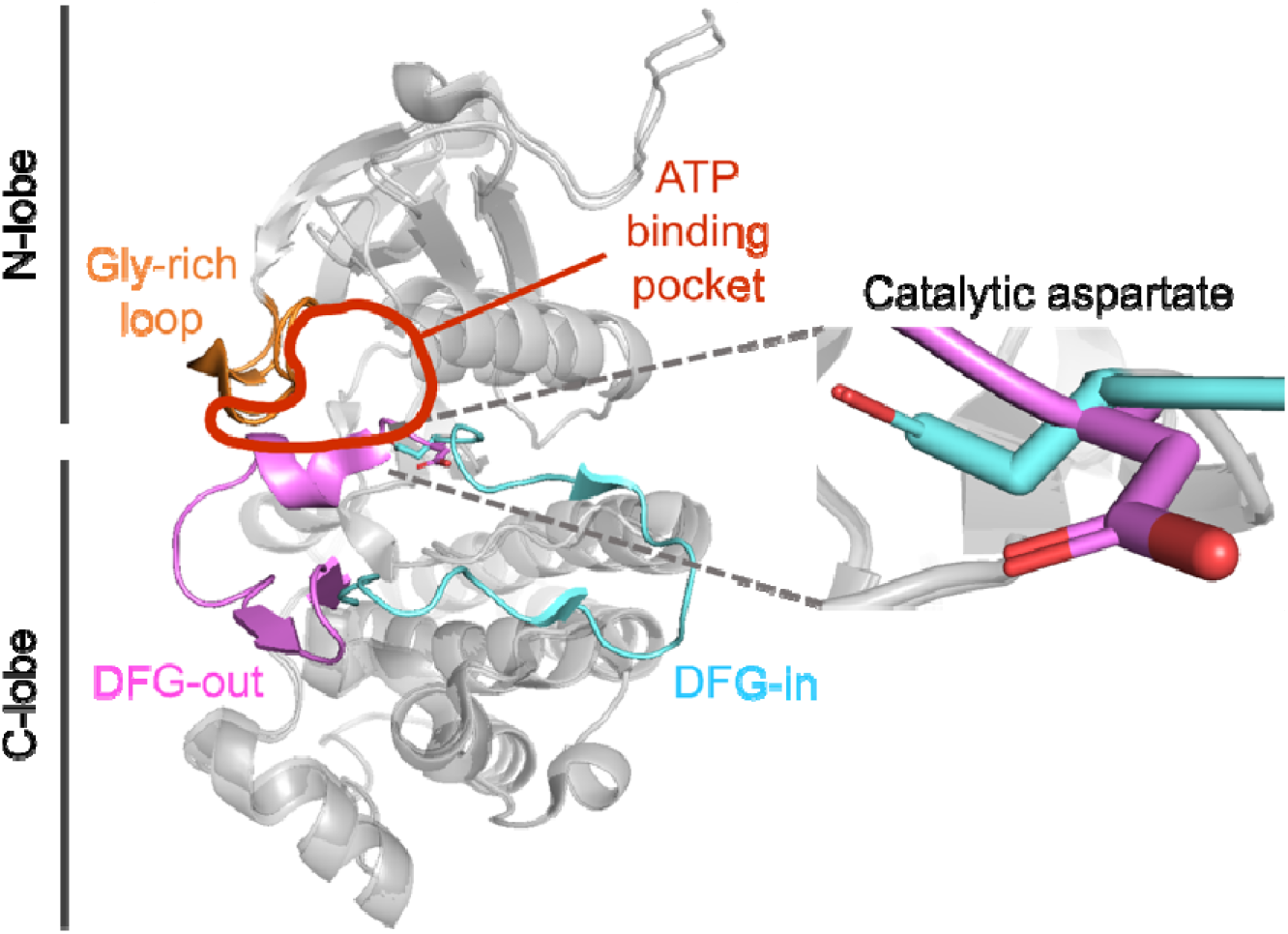
A representative protein kinase structure in DFG-in/-out states. DFG-in and DFG-out conformations of Abl kinase were superposed (PDB entries 3KF4 and 3KFA, respectively).^19^ In the DFG-in conformation (cyan),activation loop exposes the ATP binding pocket while orienting the catalytic aspartate towards the ATP bindin pocket. In the DFG-out conformation (pink), activation loop occludes the ATP binding pocket and orients the catalytic aspartate away from the ATP binding pocket. The glycine-rich loop is shown in orange.

**Figure 2.**
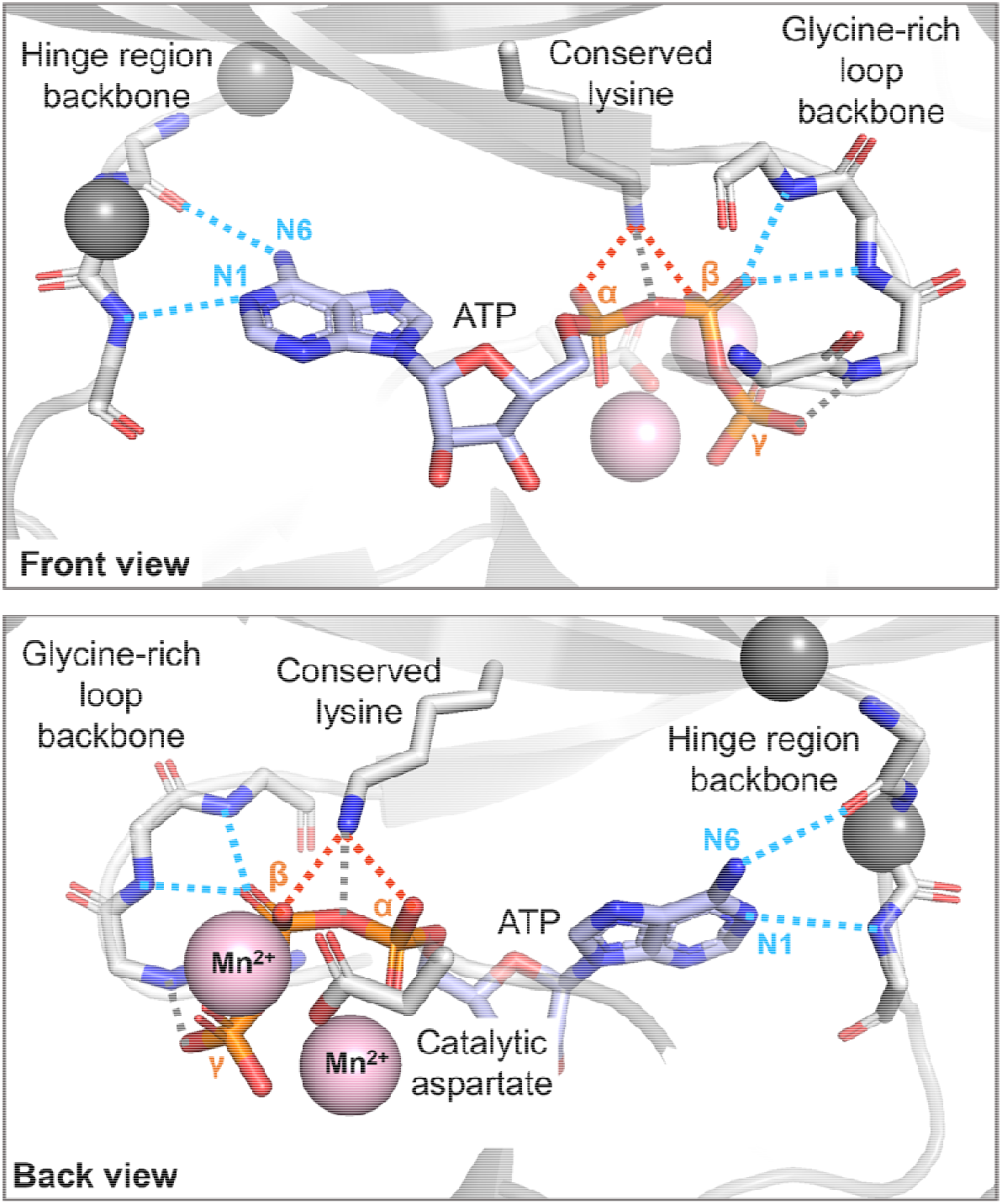
The ATP-bound catalytic subunit of cAMP-dependent protein kinase (PDB entry 1ATP).^20^ The ATP, hinge region and glycine-rich loop backbone, the conserved lysine, and the catalytic aspartate is shown in sticks. Manganese ions are depicted as pink spheres. The position of two common gatekeeper residues is shown as smaller gray spheres. The blue, red, and orange colored atoms correspond to nitrogen, oxygen, and phosphorus, respectively. The possible dipole-dipole (hydrogen bond), ion-ion (salt bridge), and ion-dipole interactions between the kinase and ATP are depicted in blue, red, and gray colored dashed lines, respectively.

### 1.3. Protein kinase deregulation due to point mutations

Kinase point mutations, occurring at the kinase’s functionally important positions, impact cell fate by altering the specificity of the enzyme or creating a constitutively active protein kinase. These changes eventually lead to severe clinical outcomes. For example, naturally occurring bulkier side chain substitutions at the gatekeeper residue cause drug resistance, as this change blocks the entry of ATP binding. Two such gatekeeper mutation examples are T315I substitution in ABL kinase and T790M substitution in EGFR. Constitutive activity of these mutants is attributed to the interaction between the phenylalanine of the DFG motif and the mutant gate-keeper residue^21–25^ There are also frequent mutations found within the N-lobe side of the activation loop.^26^ For example, V600 of BRAF is a commonly mutated residue in cancer. The V600D/E/G/K/L/M/R mutations of BRAF protein kinase were shown to lead to an over-active kinase. This positional change corresponds to the activating mutations D1228V/N/H in MET and D816E/H/V/N/F/Y/I in KIT. Another prominent example is the L858R (activation loop) and G719S (proximal to glycine-rich loop) mutants of EGFR, which destabilize the inactive conformation of the kinase.

### 1.3. Utilization of kinase mutations to study kinase function

ATP binding pocket point mutations have been exploited by drug discovery and chemical genetics approaches to dissect the kinase function. In the case of drug discovery, protein kinase gatekeeper residues and other critical residues are artificially mutated in model systems to study the mechanism of drug resistance. This approach allows scientists to tailor novel compounds to be utilized for the treatment of drug resistant tumors.^27^ Within the context of chemical genetics, naturally occurring mutations are used as a tool to evaluate the biological significance of a protein kinase.^28^ For this, the kinase gatekeeper residue is mutated to either glycine or alanine, which allows binding of a bulky ATP analogue. So, it provides a useful means to turn the enzyme off without imposing any genetic modification.^29,30^Some of these engineered kinases include yeast v-Src (I338G), c-Abl (T315A), Cdk2 (F80G), and Mps1 (M516G),7 and human Cdk12 (F813G) mutants.^31^ Besides their numerous advantages, exploiting ATP binding pocket mutations for different research purposes contains important risk factors, such as the improper choice of mutation-drug combinations or inactivation of the enzyme upon mutation. To minimize these risks, pre-screening of mutation-drug combinations with binding affinity prediction tools stands out as a prominent alternative.

### 1.4. How far are we in predicting the impact of kinase mutations on their ligand binding abilities?

Several methods have been proposed to predict protein-ligand binding affinities. Though, only recently, the accuracy of these methods has been evaluated within the scope of predicting the impact of kinase mutations in ligand binding. In 2018, Hauser *et al.*, compiled a set, involving 144 ligand affinity data regarding clinically diagnosed Abl mutations.^32^ A year later, in 2019, Aldeghi *et al.* assessed the capacity of statistical mechanics, mixed physics- and knowledge-based potentials, and machine learning approaches in predicting the impact of 31 Abl mutations with their drug interactions^33^. Both of these approaches used a hybrid structure-based Abl benchmark, composed of wild type co-crystal/docked structures and modeled mutant cases. Moreover, both papers assessed the use of sophisticated tools, the employment of which would be too complicated for many experimental biologists. Next to these approaches, there are also web-based protein-ligand binding affinity predictors, which can easily be used by the non-experts to plan/guide their experiments. Though, the assessment of these servers within the context of kinase mutations has never been done. To compensate for this shortcoming, we have benchmarked four web-based protein-ligand scoring functions, DSX-ONLINE, KDEEP, Pose&Rank, PRODIGY-LIG, as well as two general scoring functions, PDBePISA, HADDOCK Score^34–41^ to assess their capability in predicting the impact of kinase mutations on their ligand binding. We selected these predictors based on their user-friendliness and their widespread use. The benchmarking has been performed on our structure-based **BINDKIN** (effect of point mutations on the **BIND**ing affinity of protein **KIN**ase-ligand complexes) data set.

Our BINDKIN benchmark is composed of 23 experimentally determined wild type and mutant kinase structures co-crystallized with their ligands. It covers nine kinases (seven EGFR, three Abl, three Mps1, three Src, two Cdk2, one ALK, one FGFR, one Kit, and one PKA), 15 unique point mutations and binding modes of 18 different ligands. The affinity data associated to each benchmark case is reported in the form of IC_50_, K_d_, and K_i_ (Table S1). Majority of the presented mutation positions are within or in the vicinity of the ATP binding pocket. The web-server performances over BINDKIN show that the raw server scores fail to produce good correlations with the experimental data. However, when we started looking in to the direction of change (whether a mutation improves/worsens the binding), we observe that for K_i_, DSX-ONLINE could predict the impact of point mutations on the binding affinity accurately, while achieving a Pearson’s R value as 0.97 (n=6). The other kinetic metrics lead to less sensitive mutation-induced changes, thus could not be correlated with any of the scoring terms. Expanding on the DSX’s success in reproducing K_i_ changes, we have probed DSX-ONLINE also on the BINDKIN-homology benchmark, which contains the homology models of our K_i_ cases. This resulted in a drop of R from 0.97 to 0.45, highlighting the importance of structure quality in the rapid prediction of kinase mutations’ impact. We hope that these results will guide the experimentalists in designing their kinase mutation experiments, and the computational biologists in improving their protein-ligand scoring functions. The BINDKIN results, server outputs, and other related results are publicly available at https://github.com/CSB-KaracaLab/BINDKIN

## 2. METHODS

### 2.1. The collection of the BINDKIN benchmark

To construct the **BINDKIN** (effect of point mutations on the **BIND**ing affinity of protein **KIN**ase-ligand complexes) benchmark, we performed a thorough search in the Protein Data Bank (PDB)^42^ (https://www.rcsb.org/) and obtained the list of available wild type and mutant kinase-ligand complexes. Our final list was obtained by imposing the following criteria:

- For each mutant complex, there has to be a wild type complex, containing the same protein and the ligand.
- The wild type and mutant complexes should be determined in the same study (i.e., they should come from the same paper).
- For each complex, there has to be experimentally determined binding affinity available in the form of IC_50_, K_d_, or K_i_ together with a related research paper. The experimental binding kinetics data were acquired from PDBbind (http://www.pdbbind-cn.org/index.php),^43–45^ Binding DB (https://www.bindingdb.org/bind/index.jsp),^46^ and Binding MOAD (http://bindingmoad.org/Search/advancesearch)^47,48^databases.
- The ligand has to be a non-covalent one.

This limitation has left us with 23 wild type-mutant complex pairs, making up the BINDKIN benchmark. BINDKIN constitutes of seven EGFR, three Abl, three Mps1, three Src, two Cdk2, one ALK, one FGFR, one Kit, and one PKA proteins. These complexes present 15 unique point mutations distributed. BINDKIN contains binding modes of 18 different ligands. The pharmacophoric characteristics of the ligands were evaluated with the ALOGPS web server of the Virtual Computational Chemistry Laboratory (http://www.virtuallaboratory.org/web/alogps/)^49–58^ and pkCSM web server (http://structure.bioc.cam.ac.uk/pkcsm).^59^ The available ligand-related data were obtained from RCSB PDB and PubChem (https://pubchem.ncbi.nlm.nih.gov/). Most of the mutation positions are within or in the vicinity of the ATP binding pocket (Figure 3A). All the benchmark related features are deposited in https://github.com/CSB-KaracaLab/BINDKIN. Further details about the benchmark characteristics are described in the Results section.

**Figure 3.**
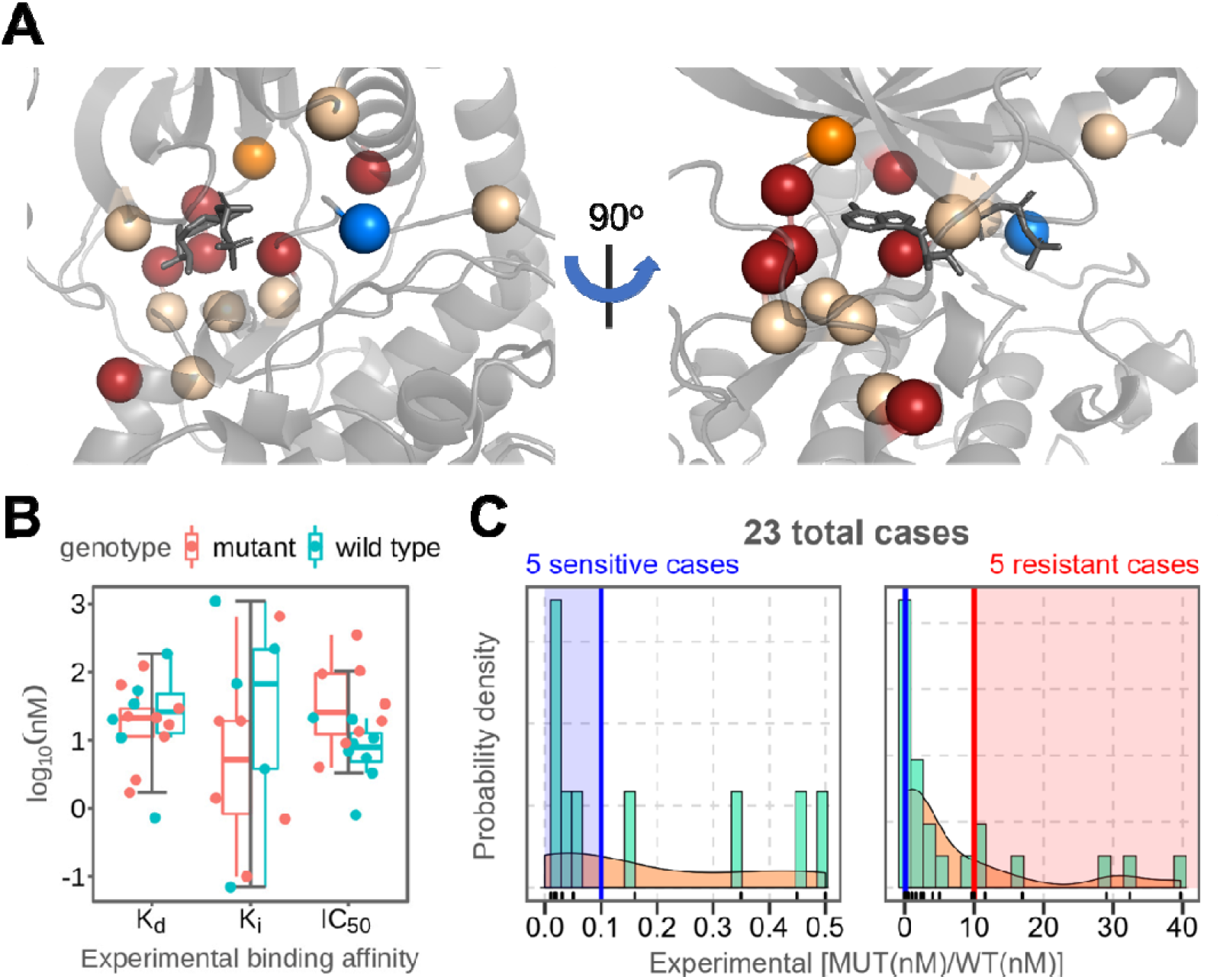
The BINDKIN benchmark characteristics. **(A)** The corresponding positions of the mutated residues are mapped on a representative ATP-bound protein kinase (cAMP-dependent protein kinase with PDB entry 1ATP).^20^ The orange sphere is the position where the majority of the BINDKIN mutations are located. The blue sphere indicates the *sensitizing* mutations positions, where red spheres indicate the mutations conferring *resistance*, an light brown ones indicate unclassified mutations. **(B)** Distribution of the experimental binding kinetics data for the 42 individual cases. The data was split as K_d_, K_i_, and IC_50_ and plotted as box-and-whisker plots for the corresponding wild type and mutant cases. **(C)** Probability distribution of the experimental kinetic data for wild typ and mutant pairs. The blue and red areas indicate the *drug-sensitive* and *drug-resistant* cases, respectively. The KDE (kernel density estimate) is represented as the light-brown area.

### 2.2. The BINDKIN-homology benchmark

The wild type and mutant sequences were structurally modeled with the default settings of the I-TASSER web server (https://zhanglab.ccmb.med.umich.edu/I-TASSER/).^60–62^ During modeling, the original coordinates of the wild type or mutant structures were excluded from the template list. After obtaining the homology models, their corresponding ligands were placed at their catalytic binding pocket by using the original crystal structure as a template. The fitting was performed with PyMOL.^63^ In these crude models, steric clashes were observed. To optimize the protein-ligand interface, we applied water refinement on each model by using the HADDOCK2.2 refinement interface (https://milou.science.uu.nl/services/HADDOCK2.2/haddockserver-refinement.html).^38^

### 2.3. The benchmarked web servers

Before benchmarking, the crystallization buffer additives, ions were discarded from the co-crystal structures (in the .pdb format). For each occurrence of multiple conformations, the conformer with the highest occupancy was retained and the other conformers were removed. In one of the EGFR cases (5em7), the ligand 5Q4 is depicted in two different conformations. Both conformations were taken into consideration. The ligand coordinate files were converted to ‘mol2’ and ‘sdf’ formats based on the different file format requirements of the web servers. For mol2 conversion Open BABEL (v2.4.1) and for sdf conversion the Online SMILES Translator and Structure File Generator (National Cancer Institute) (https://cactus.nci.nih.gov/translate/) were used. For the sdf conversion, the “Aromatic” SMILES representation option (prints the unique SMILES string into the ‘sdf’ file) and “3D” coordinates option were chosen. The sdf files were protonated by default. The conformations and the 3D coordinates of the ligands were retained during conversion to both mol2 and sdf formats.

Six different web-based scoring functions were used to predict the binding affinities listed within BINDKIN. A short description of each web server is provided in Table 1.

**Table 1:**
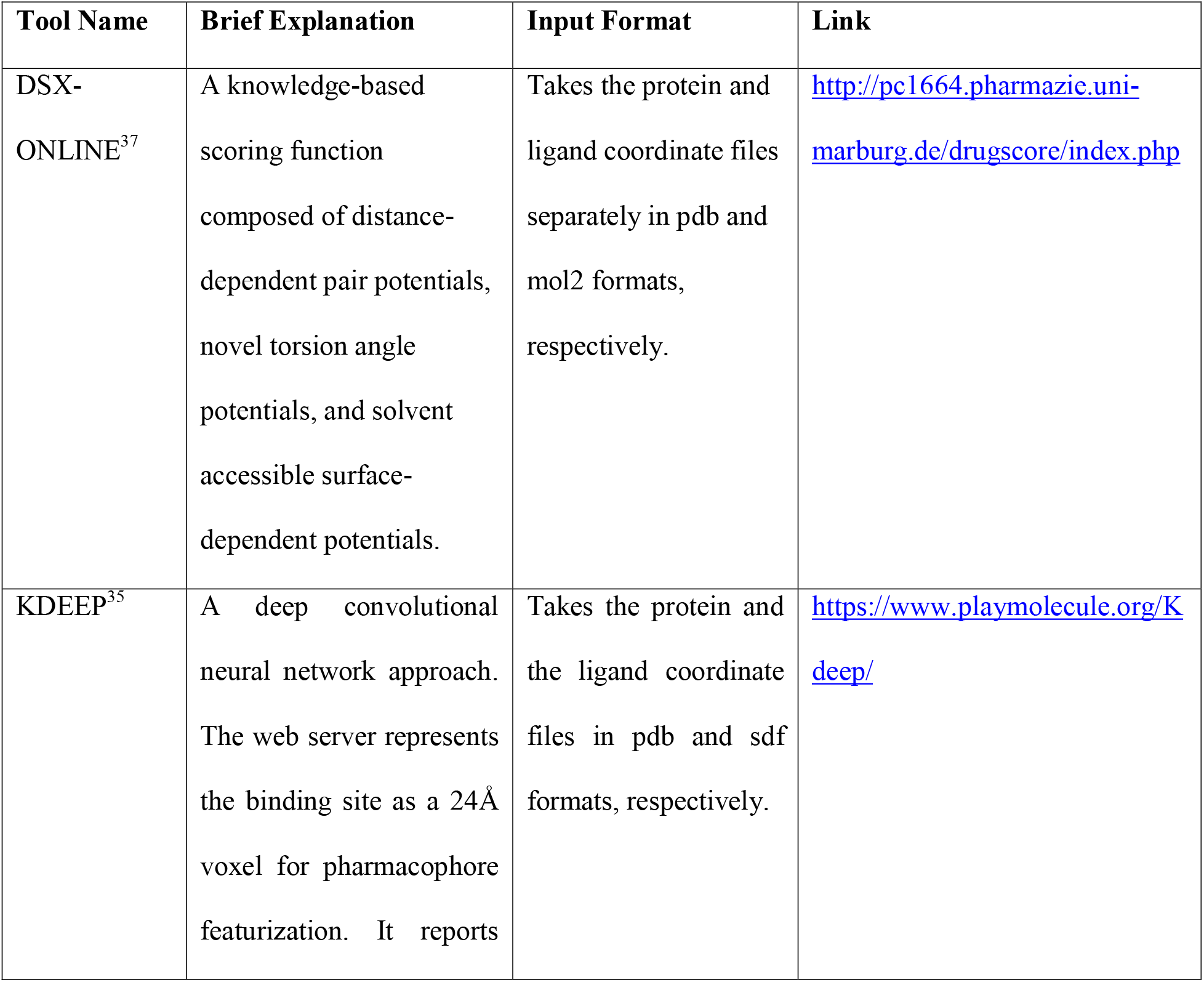
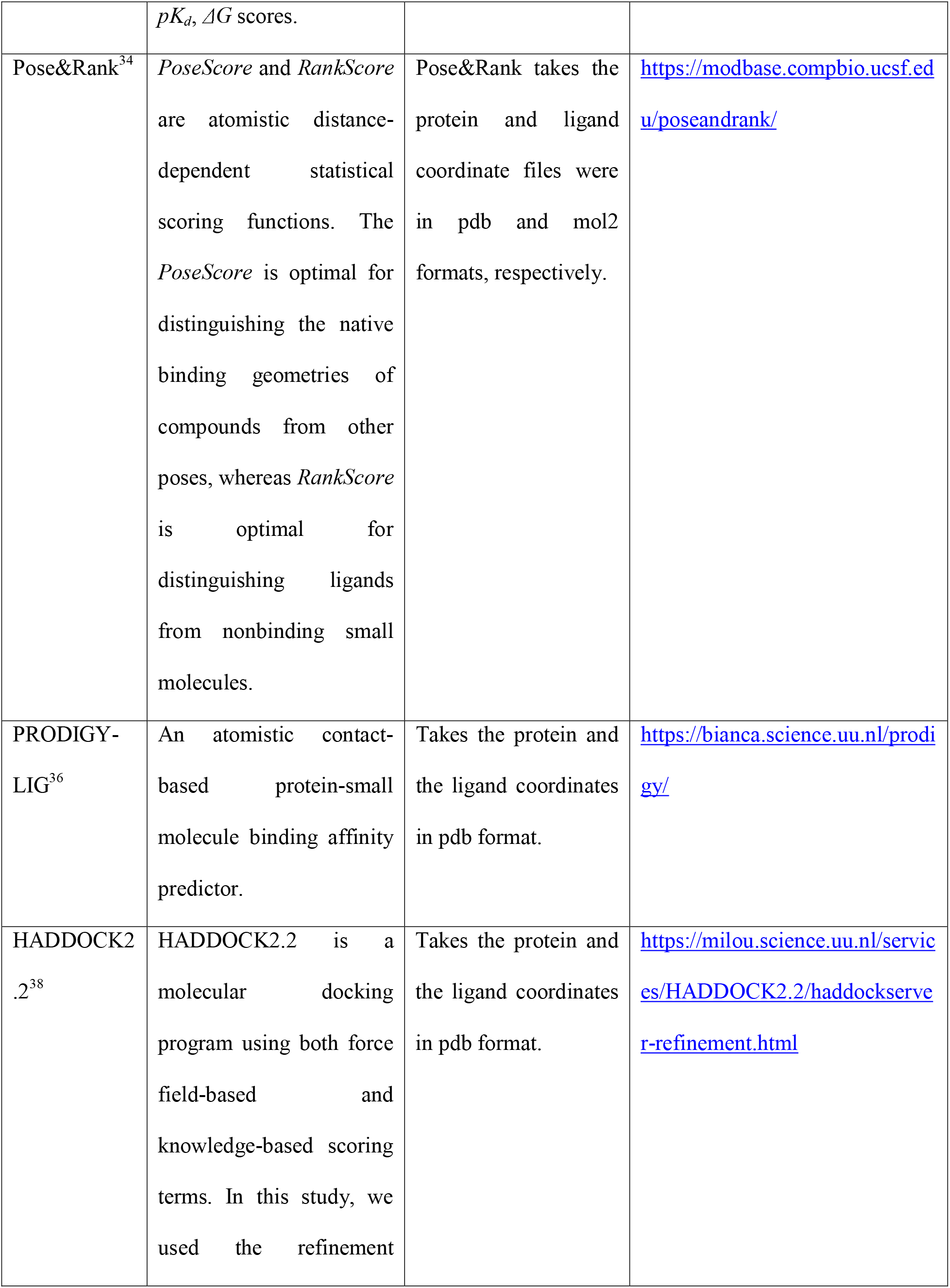
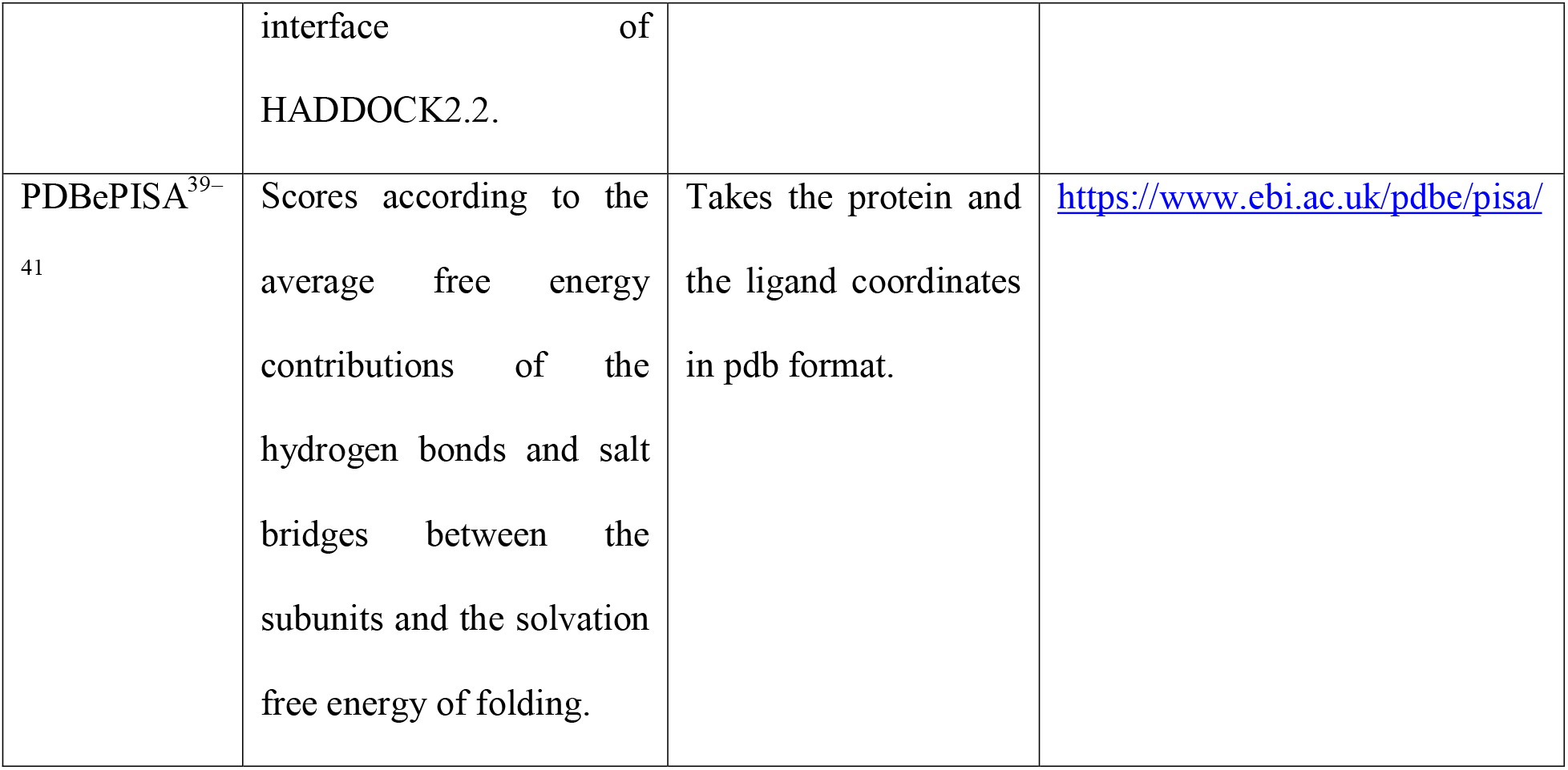
The list of the benchmarked servers.

### 2.4. The assessment of the prediction results

Three different approaches were used to assess the prediction results. These are *direct assessment*, *delta assessment*, and *binary assessment*. *Direct assessment* is the classical approach in which the raw scores were correlated with the experimental ones by linear regression. In the *delta assessment*, mutation-induced affinity (1) and score (2) changes were correlated.

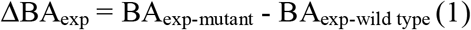

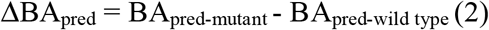

In the *binary assessment*, we checked the agreement in the mutation induced direction change. In this case, if a mutation leads to a decrease in the experimental binding affinity and it is predicted as such by the computational prediction, it was considered as a successful prediction. If the increase in the experimental affinity is accompanied by a decrease in the predicted affinity, then this case was counted as a misprediction. In the *binary assessment*, the percentage of correct predictions was used as the final success rate.

For the linear regression analyses, the associated R and p values, and the plots were generated with R (v3.6.1)^64^, invoked by RStudio (v1.2.5001).^65^ The following R packages were used to generate scatterplots, barplots, or histograms: ggpubr,^66^ ggplot2,^67^ magrittr,^68^ ggrepel,^69^ grid,^64^ and gridExtra.^70^ The heatmaps were produced with pheatmap^71^ and RColorBrewer^72^ R packages.

## 3. RESULTS AND DISCUSSION

### 3.1. The BINDKIN benchmark characteristics

Our benchmark was gathered from the literature by following the criteria listed under Methods. BINDKIN benchmark covers nine protein kinases (seven EGFR, three Abl, three Mps1, three Src, two Cdk2, one ALK, one FGFR, one Kit, and one PKA), where each wild type kinase has a mutant counterpart, leading to 23 pairs (with 8, 9, and 6; IC_50_-, K_d_-, and K_i_-associated cases, respectively) and 42 individual cases (with 16, 15, and 11; IC_50_-, K_d_-, and K_i_-cases, respectively) (Table S1). It includes 15 different mutations, corresponding mostly to single point ones (except one quadrupole, two triple, and two double mutations). The mutations are distributed within or in the vicinity of ATP binding pocket (Figure 3A). In their wild type states, these mutation positions exert diverse biophysical characteristics (charged, polar, hydrophobic), whereas in their mutant forms they predominantly turn into hydrophobic and charged amino acids. Also, half of the mutations result in bulkier residues. The 18 different ligands presented in BINDKIN are ATP-competing chemicals. They are nitrogen-rich and mainly hydrogen bond acceptors. From the degrees of freedom point of view, BINDKIN ligands cover a diverse range of rotatable bonds, i.e., from 0 to 11.

The binding affinity range spanned by BINDKIN is 0.80 - 350, 0.73 - 185, and 0.07 - 1090 nM for IC_50_, K_d_, and K_i_ subsets, respectively (Table S2).^73–88^ So, the K_i_ subset covers the widest affinity range (Figure 3B). We classified a mutation as *sensitive* if the mutant complex's binding affinity is at least ten-fold higher than that of its wild type state.^32,33,89^ When the mutant complex's binding affinity is less than one-tenth of its wild type, then this case is classified as *resistant*. According to this criterion, BINDKIN contains five *sensitive* (four K_i_ and one K_d_) and five *resistant* cases (three IC_50_, and two K_d_) (Figure 3C). The probability density of affinity values is left-skewed, meaning that our benchmark is biased towards *drug-sensitizing* mutations.

### 3.2. Majority of the web-based scoring functions predicts the direction of affinity change caused upon mutation

Advanced computational tools have been shown to accurately predict protein kinase-ligand binding affinities.^33,89^Considering that these approaches do not have a broad reach among the experimental biology community, in this work, we have focused on user-friendly web servers' performance within the context of predicting the impact of kinase mutations. To this end, we have tested the prediction performances of DSX-ONLINE, HADDOCK2.2 (refinement interface), KDEEP, PDBePISA, Pose&Rank, and PRODIGY-LIG over BINDKIN.

To assess the performance of each tool, we have pooled the affinity predictions (n=42) and analyzed their association with the experimental data. This *direct assessment* shows that the presented scoring functions could not predict the affinity of BINDKIN cases (Figure 4). Here, the highest R-value is produced by KDEEP (R=0.31). When we divide the same set according to their kinetic metrics (IC_50_, K_d_, K_i_), the highest correlation is obtained with Rank and PRODIGY-LIG scores over IC_50_ cases (R=0.55). Considering wild type and mutant states separately did not improve the presented correlations (data not shown).

**Figure 4.**
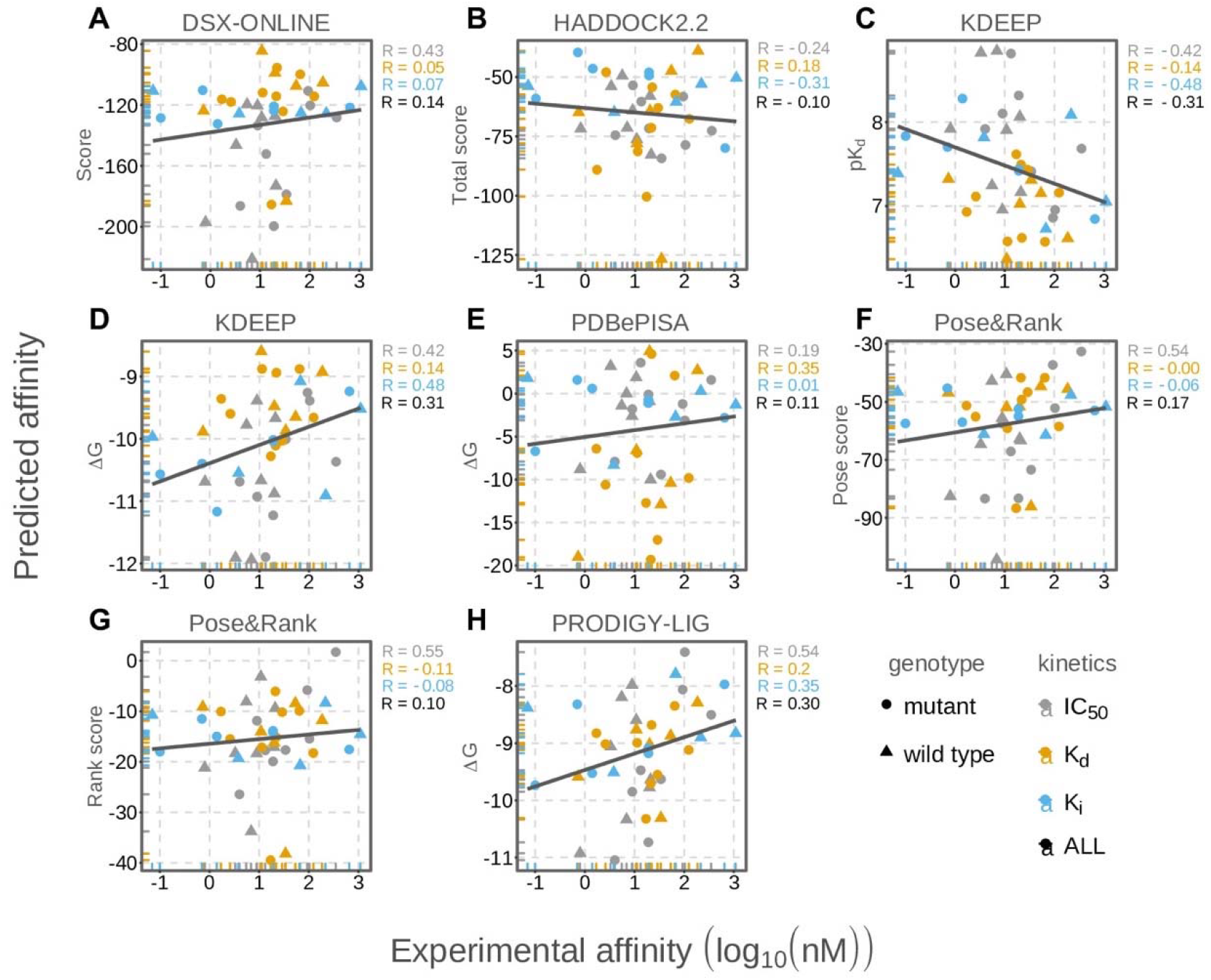
The performance of the various scoring functions accorring to *direct assessment*. Due to the nature of the prediction scores, a negative correlation for pKd (KDEEP) and a positive correlation for rest of the tools is expected. **(A)** DSX-ONLINE, **(B)** HADDOCK2.2 (*HADDOCK score*), **(C, D)** KDEEP (*pK_d_* and *ΔG*), **(E)** PDBePISA (*ΔG*), **(F, G)** Pose&Rank (*PoseScore* and *RankScore*), and **(H)** PRODIGY-LIG (*ΔG*). The sample size for IC_50_, K_d_, and K_i_ are n=16, n=15, and n=11, respectively.

Next to the *direct assessment*, we have also performed a relative evaluation of the results with *delta* and *binary assessments* (Figure 5, see Methods). *Delta assessment* measures the absolute change in the (predicted) affinity upon mutation. Using *delta assessment* improves R values significantly. This is especially the case for the R-performances of KDEEP and DSX-ONLINE over K_i_-cases, which become 0.75 and 0.97, respectively. In the *binary assessment*, the accuracy of predictors was measured by analyzing the agreement between the direction of change in the experimental and predicted binding affinities. According to this metric, DSX-ONLINE accurately predicts the affinity change of all K_i_-cases. The success rate of DSX-ONLINE over K_i_-cases is followed by *PoseScore* and PRODIGY-LIG, both leading to 83% accuracy. As another important highlight, KDEEP (*pK_d_*) reaches 88% success rate on IC_50_ cases. As an across-metric comparison, over K_i_-cases, DSX-ONLINE leads to an R of 0.07 and 0.97 in *direct* and *delta assessments*, and 100% accuracy in *binary assessment*. While *delta* and *binary assessments* agree in this particular case, in some other cases, they contradict. As an example, PRODIGY-LIG achieves an R value of 0.29 in the *delta assessment*, and 83% accuracy in the *binary assessment*. The contradiction between these two types of assessment approaches can be explained by the data spread. For example, a scoring function may achieve a great success on the *binary assessment*, because it can predict the direction of affinity change. However, the same scoring function may not be able to predict the amount of change, which would lead to a lower R value. Within the context of the biological meaning of the assessment approaches, we suggest the concurrent use of *delta* and *binary assessments*.

**Figure 5.**
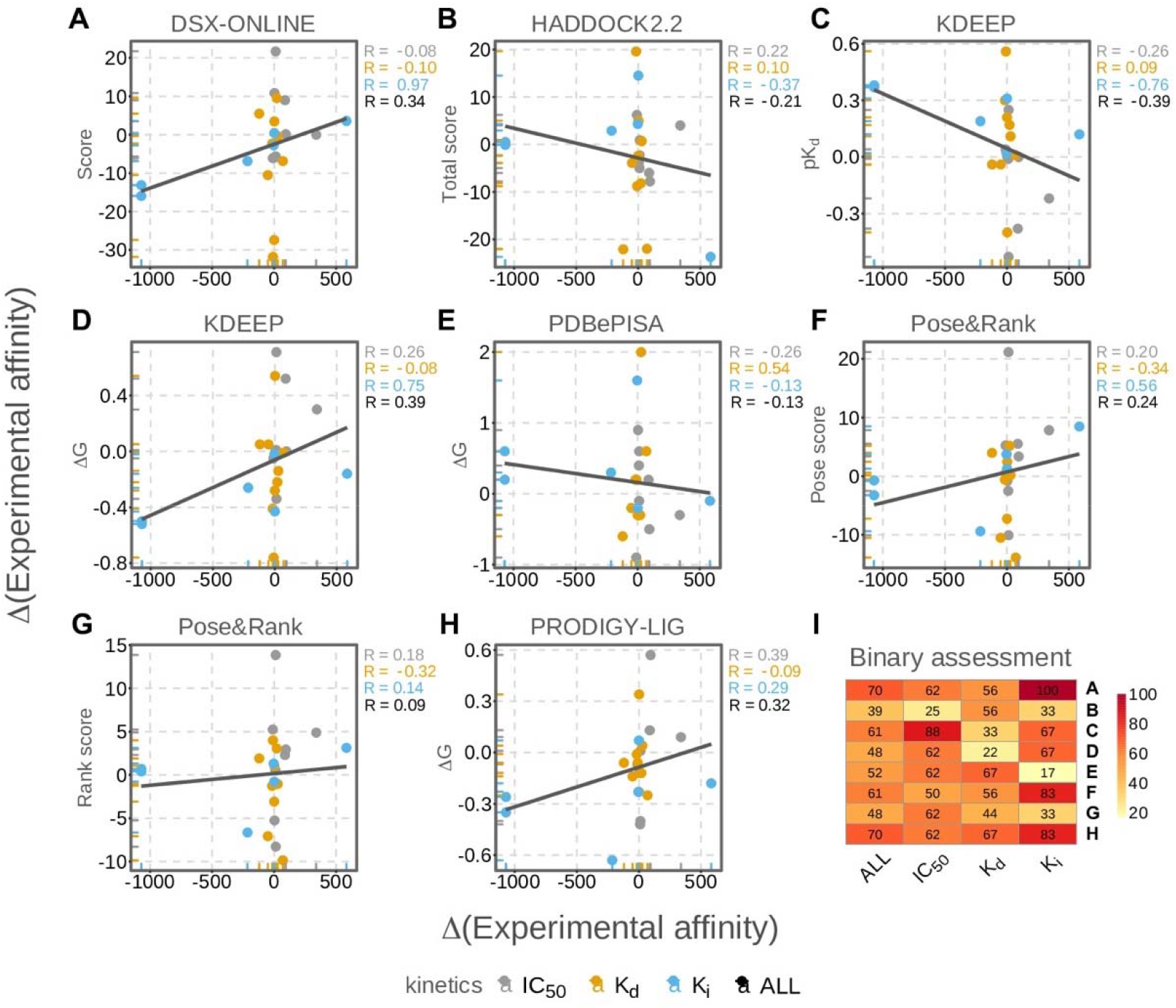
The performance of the various scoring functions according to the *delta assessment*. Due to the nature of the prediction scores, a negative correlation for pKd (KDEEP) and a positive correlation for rest of the tools is expected. **(A)** DSX-ONLINE, **(B)** HADDOCK2.2 (*HADDOCK score*), **(C, D)** KDEEP (*pK*_*d*_ and *ΔG*), **(E)** DBePISA (*ΔG*), **(F, G)** Pose&Rank (*PoseScore* and *RankScore*), and **(H)** PRODIGY-LIG (*ΔG*). **(I)** The success rates from the *binary assessment* are provided as a heatmap. Yellow indicates low accuracy and red indicates high accuracy predictions. The sample size for IC_50_, K_d_, and K_i_ are n=8, n=9, and n=6, respectively.

Among the three different kinetic data types, according to the *direct assessment*, IC_50_ turns out to be the best descriptor for most of the scoring functions. When the complexes are considered as mutant/wild type pairs (*delta* and *binary assessments*), K_i_ comes out as the best descriptor (Figure 5). Interestingly, none of the methods could lead to a reliable prediction over the K_d_ set. K_d_ is a universal measure of binding while K_i_ is defined for enzymes only, which could be the reason of the high R values reported for the K_i_-set.^90^ However, we also cannot rule out the fact that success rates of the kinetic descriptors may depend on the number of cases present for each descriptor (8 pairs for IC_50_, 9 pairs for K_d_, and 6 pairs for K_i_).

In a real case scenario, the structure of the complex under study might not be available. Expanding on this, we have homology modeled the K_i_-cases and probed these structures with DSX-ONLINE (as this combination gives the best *delta and binary assessments*). The mutation-specific features such as hydropathy index change^91^, residue volume change^92^, and drug resistance or sensitivity were chosen for the characterization of predicted affinities (Figure 6). As a result of this exercise, here, we show that the performance of DSX-ONLINE is dramatically worsened by the worsened quality of the input structure (an R-drop from 0.97 to 0.45). We also observe that a decrease in the residue volume upon mutation is accompanied by an increase of polarity and drug sensitivity. However, in this case, the amount of cases is not sufficient to draw a conclusion regarding the effect of mutation characteristics on the prediction accuracy.

**Figure 6.**
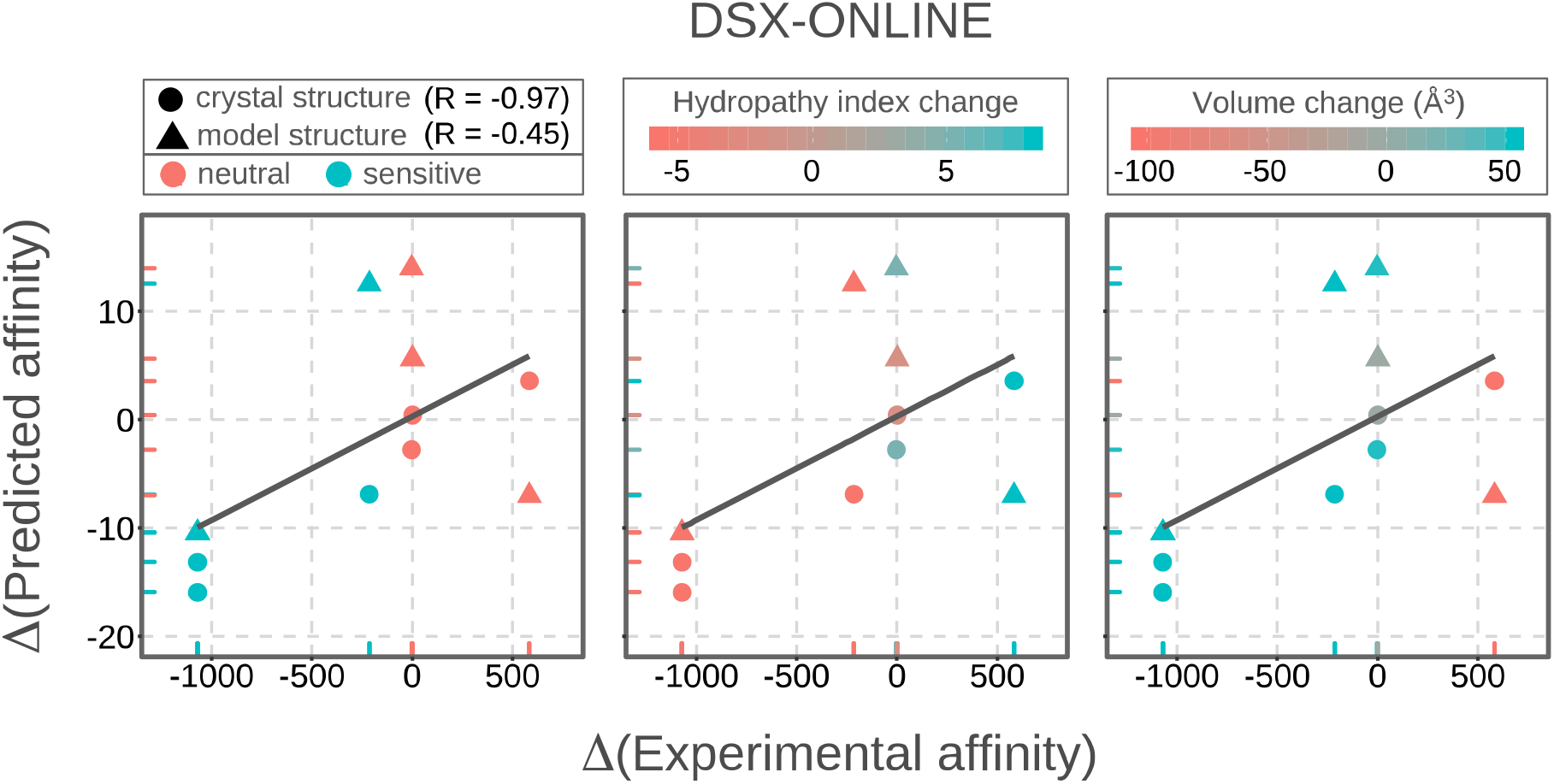
The default scoring function of DSX-ONLINE was tested on the K_i_-associated crystal structure and homology model structure sets by using *delta assessment*. Orange color corresponds to the neutral cases (left) or a decrease in the hydropathy index (middle) or residue volume (right). Turquoise color corresponds to the sensitive cases (left) or an increase in the hydropathy index (middle) or residue volume (right). RMSD: 2.0 ± 0.5. The sample size of the crystal structures and model structures are n=6 and n=5, respectively.

## 4. CONCLUSIONS

The presented study investigates whether web-based scoring functions can predict the impact of kinase mutations. To test this, we have compiled BINDKIN, the first structure-based affinity benchmark of wild type and mutant protein kinase-small molecule complexes. Based on the K_i_-associated cases of BINDKIN, we have also compiled a homology model benchmark. Using these structure sets and their available experimentally binding affinity data, we have assessed the prediction accuracy of six different web servers. These are DSX-ONLINE, HADDOCK2.2, KDEEP, PDBePISA, Pose&Rank, and PRODIGY-LIG. The prediction accuracy of the web servers has been thoroughly analyzed by the use of different assessment approaches. As a result, we have observed that (i) K_i_ is a sensitive descriptor that can capture the effect of mutations on binding affinity, (ii) DSX-ONLINE is the best tool in predicting the mutation-induced K_i_ changes, (iii) the quality of the input structure directly impacts the performance of DSX-ONLINE, and (iv) more kinase mutation-related ligand binding data are needed to perform a statistically significant analysis.

## ASSOCIATED CONTENT

Supplementary Information and Supplementary Tables are submitted as a separate file.

## AUTHOR CONTRIBUTIONS

This research was devised and performed by ME, TK, and EK. The paper was written by ME and EK. All authors contributed to careful revising of the manuscript.

## NOTES

The authors declare no competing financial interest.

## ACKNOWLEDGEMENTS

We thank Dr. Gerard Martinez (Acellera Ltd., London, United Kingdom) for his help and guidance during benchmarking of the KDEEP web server. We thank Dr. Daniel Fisher (The Institute of Molecular Genetics of Montpellier (IGMM), Montpellier, France) for helping us clarifying the wrong binding kinetics data records found in the PDBbind database (PDB entries 4EOR and 4EOK).

