## Supplementary for "How far are we in the rapid prediction of drug resistance caused by kinase mutations?"

**Supplementary Information**

**The proposed catalytic mechanism and kinetics for the protein kinases**

The catalytic aspartate is responsible for transfer of the γ-phosphate from ATP to the target residue of a substrate protein. According to an accepted model, the ion-dipole interaction between the negatively charged oxygen of catalytic aspartate carboxylate and proton of the protein substrate hydroxyl group, lowers the pK_a_ value of the substrate hydroxyl. This results in conversion of the alcohol group of the target residue into an alkoxide at physiological pH, which is a stronger nucleophile than alcohol. Upon correct positioning of ATP and the substrate protein, nucleophilic attack of the alkoxide to the phosphorus atom of γ-phosphate initiates the enzymatic reaction. On the other hand, pH dependence studies are in conflict with this general acid catalysis model, suggesting a dissociative-like transition state. As in the case of explained catalytic models, kinetic models of protein kinases are also in conflict. The reactions catalyzed by protein kinases require both ATP and a substrate protein, thus, can be viewed as bisubstrate kinetic mechanisms. Bisubstrate reaction kinetics with compulsory ordered ternary complex mechanism was experimentally observed for PI3Kα and p38 MAPKα.^1,2^ On the other hand, these results vary by use of different peptide substrates. Also, the steady-state kinetic approaches have revealed that several important protein kinases have shown random bisubstrate kinetic mechanism, again including p38 MAPKα.^3–7^

**Supplementary Tables**

**Table S1.** The content of BINDKIN. The respective PDB entries, protein names, mutation remarks, the experimental binding kinetics data, and references are provided in different columns. The repeating entries are highlighted in bold under the Remarks section.

| **PDB entry** | **Protein Kinase** | **Remarks** | **Affinity data type** | **Affinity data (nM)** | **References** |
| --- | --- | --- | --- | --- | --- |
| 2qoh | ABL1 | **WT** | IC_50_ | 20 | ^8^ |
| 2z60 | ABL1 | **MUT (T315I)** | IC_50_ | 9 | ^8^ |
| 2iw9 | Cdk2 | WT | IC_50_ | 8.9 | ^9^ |
| 2iw8 | Cdk2 | MUT (F82H, L83V, H84D) | IC_50_ | 103 | ^9^ |
| 3f3v | Src | **WT** | IC_50_ | 21 | ^10^ |
| 3f3w | Src | **MUT (T338M)** | IC_50_ | 34 | ^10^ |
| 3qri | ABL1 | **WT** | IC_50_ | 0.8 | ^11^ |
| 3qrj | ABL1 | **MUT (T315I)** | IC_50_ | 4 | ^11^ |
| 3w2s | EGFR | **WT** | IC_50_ | 6.9 | ^12^ |
| 3w2r | EGFR | **MUT (T790M, L858R)** | IC_50_ | 19 | ^12^ |
| 5ap0 | MPS1 (TTK) | **WT** | IC_50_ | 5.5 | ^13^ |
| 5ap3 | MPS1 (TTK) | **MUT (S611G)** | IC_50_ | 93 | ^13^ |
| 5ap1 | MPS1 (TTK) | **WT** | IC_50_ | 10.8 | ^13^ |
| 5ap4 | MPS1 (TTK) | **MUT (S611G)** | IC_50_ | 350 | ^13^ |
| 5ap2 | MPS1 (TTK) | WT | IC_50_ | 3.3 | ^13^ |
| 5ap6 | MPS1 (TTK) | MUT (C604W) | IC_50_ | 13.3 | ^13^ |
| 1xh4 | PKA | WT | K_d_ | 34 | ^14^ |
| 1xh9 | PKA | MUT (Q84E, V123A, L173M, Q181K, F187L) | K_d_ | 17 | ^14^ |
| 2ity | EGFR | **WT** | K_d_ | 53.5 | ^15^ |
| 2ito | EGFR | **MUT (G719S)** | K_d_ | 123.6 | ^15^ |
| 2itz | EGFR | **MUT (L858R)** | K_d_ | 2.6 | ^15^ |
| 2j6m | EGFR | **WT** | K_d_ | 10.9 | ^15^ |
| 2itp | EGFR | **MUT (G719S)** | K_d_ | 11.3 | ^15^ |
| 2itt | EGFR | **MUT (L858R)** | K_d_ | 1.7 | ^15^ |
| 3g0e | KIT | WT | K_d_ | 20 | ^16^ |
| 3g0f | KIT | MUT (D816H) | K_d_ | 22 | ^16^ |
| 4mxo | Src | **WT** | K_d_ | 0.73 | ^17^ |
| 4mxx | Src | MUT (A403T) | K_d_ | 29 | ^17^ |
| 4mxz | Src | **MUT (T338M)** | K_d_ | 21.2 | ^17^ |
| 5am6 | FGFR | WT | K_d_ | 185 | ^18^ |
| 5am7 | FGFR | MUT (V561M) | K_d_ | 64.5 | ^18^ |
| 4cli | ALK | WT | K_i_ | 0.07 | ^19^ |
| 4clj | ALK | MUT (L1196M) | K_i_ | 0.7 | ^19^ |
| 4eor | Cdk2 | WT | K_i_ | 67 | ^20^ |
| 4eok | Cdk2 | MUT (H85S, Q86M, K89D) | K_i_ | 650 | ^20^ |
| 4wa9 | ABL1 | **WT** | K_i_ | 3.8 | ^21^ |
| 4twp | ABL1 | **MUT (T315I)** | K_i_ | 0.1 | ^21^ |
| 5cav | EGFR | **WT** | K_i_ | 216 | ^22^ |
| 5cas | EGFR | **MUT (T790M, L858R)** | K_i_ | 1.4 | ^22^ |
| 5em8 | EGFR | **WT** | K_i_ | 1090 | ^23^ |
| 5em7 | EGFR | **MUT (T790M, L858R)** | K_i_ | 19 | ^23^ |

**Table S2.** The basic characteristics of BINDKIN. The number of individual cases or WT-MUT case pairs per different types of the experimental binding kinetics data (IC_50_, K_d_, and K_i_) are given. The nM concentration ranges of IC_50_, K_d_, and K_i_ subsets are given.

|  | **IC_50_** | **K_d_** | **K_i_** | **Total** |
| --- | --- | --- | --- | --- |
| #individual structures | 16 | 15 | 11 | 42 |
| #WT-MUT pairs | 8 | 9 | 6 | 23 |
| nM range | 0.8 - 350 | 0.73 - 185 | <0.07 - 1090 | <0.07 - 1090 |
